# Attention reinforces human corticofugal system to aid speech perception in noise

**DOI:** 10.1101/2020.10.22.351494

**Authors:** Caitlin N. Price, Gavin M. Bidelman

**Author notes:** **Address for editorial correspondence:** Caitlin N. Price, AuD, PhD, School of Communication Sciences & Disorders, University of Memphis, 4055 North Park Loop, Memphis, TN, 38152. **Author contributions: Caitlin N. Price:** Conceptualization, methodology, investigation, formal analysis, visualization, writing – original draft and review & editing. **Gavin M. Bidelman:** Conceptualization, methodology, formal analysis, writing – original draft and review & editing, supervision.

## Abstract

Perceiving speech-in-noise (SIN) demands precise neural coding between brainstem and cortical levels of the hearing system. Attentional processes can then select and prioritize task-relevant cues over competing background noise for successful speech perception. In animal models, brainstem-cortical interplay is achieved via descending corticofugal projections from cortex that shape midbrain responses to behaviorally-relevant sounds. Attentional engagement of corticofugal feedback may assist SIN understanding but has never been confirmed and remains highly controversial in humans. To resolve these issues, we recorded source-level, anatomically constrained brainstem frequency-following responses (FFRs) and cortical event-related potentials (ERPs) to speech via high-density EEG while listeners performed rapid SIN identification tasks. We varied attention with active vs. passive listening scenarios whereas task difficulty was manipulated with additive noise interference. Active listening (but not arousal-control tasks) exaggerated both ERPs and FFRs, confirming attentional gain extends to lower subcortical levels of speech processing. We used functional connectivity to measure the directed strength of coupling between levels and characterize “bottom-up” vs. “top-down” (corticofugal) signaling within the auditory brainstem-cortical pathway. While attention strengthened connectivity bidirectionally, corticofugal transmission disengaged under passive (but not active) SIN listening. Our findings (i) show attention enhances the brain’s transcription of speech even prior to cortex and (ii) establish a direct role of the human corticofugal feedback system as an aid to cocktail party speech perception.

**Ethics statement:** All participants provided written informed consent prior in accordance with protocols approved by the University of Memphis IRB.

**Declaration of interest:** none

## 1. Introduction

Attention is a top-down cognitive process that alerts and orients listeners to focus concentration on environmental stimuli (Petersen and Posner, 2012). In complex listening environments, attention aids the selection of behaviorally-relevant inputs over irrelevant background noise to prioritize target cues for robust speech-in-noise (SIN) understanding. Given that top-down mechanisms fine-tune auditory neural coding (Atiani et al., 2009; Gao and Suga, 2000; Suga and Ma, 2003), attention is thought to influence all stages of auditory processing from the inner ear to cortex (Galbraith et al., 2003; Hernandez-Peon, 1966; Lukas, 1980; Picton and Hillyard, 1974; Rinne et al., 2008).

Attentional modulation of auditory cortical activity is well documented (Picton and Hillyard, 1974), but whether similar enhancements exist at earlier stages prior to cortex (e.g., brainstem) remains contentiously debated. Human brainstem responses to nonspeech stimuli are largely invariant to attentional state (Hirschhorn and Michie, 1990; Picton and Hillyard, 1974; Picton et al., 1971; Woods and Hillyard, 1978). Newer contradictory findings have emerged from more recent studies on frequency-following responses (FFR)—microphonic-like potentials generated predominantly from brainstem which index neural phase-locking to dynamic sound features (Bidelman, 2018; Marsh et al., 1970). Some electrophysiological studies suggest attention enhances the robustness and temporal precision of speech FFRs (Forte et al., 2017; Galbraith et al., 2003; Hartmann and Weisz, 2019; Lehmann and Schonwiesner, 2014). Still, others demonstrate mixed (Holmes et al., 2018; Saiz-Alia et al., 2019) or even null attention-related FFR effects (Galbraith and Kane, 1993; Varghese et al., 2015). Brainstem responses are reliably recorded during sleep and sedation (Skoe and Kraus, 2010a) which bolsters long-held assumptions that subcortical processing is largely pre-attentive and automatic (Tzounopoulos and Kraus, 2009). Attention effects at the brainstem level might be too subtle to detect in scalp EEG (Varghese et al., 2015) or require highly specialized/unnatural perceptual tasks that overly challenge speech processing (Galbraith and Arroyo, 1993; Lehmann and Schonwiesner, 2014). Equivocal findings have even led to assertations that “efferent-mediated inhibitory mechanisms have [no] role to play in selective attention” and do not “suppress irrelevant information at early stages of the auditory system” (Hirschhorn and Michie, 1990, p. 507).

Architecturally, the auditory neuroaxis contains both afferent (ear-to-brain) and efferent (brain-to-ear) projections. Germane to the present experiments, descending cortico-collicular (i.e., corticofugal) fibers from primary auditory cortex (PAC) with targets in brainstem (BS) recalibrate sound processing of midbrain neurons by fine-tuning their receptive fields in response to behaviorally relevant stimuli (Suga, 2008; Suga et al., 2000). Corticofugal efferents also drive learning-induced plasticity in animals (Bajo et al., 2010; Suga et al., 2000). Given the midbrain (upper BS) is the primary source of human scalp FFRs (Bidelman, 2018; Marsh et al., 1970), these efferents may account for experience-dependent plasticity observed in seminal human FFR studies (Chandrasekaran and Kraus, 2010; Kraus and White-Schwoch, 2015; Musacchia et al., 2007; Wong et al., 2007). Despite experience-dependent changes observed in human FFRs and ample evidence for online subcortical modulation in animals (Bajo et al., 2010; Slee and David, 2015; Suga et al., 2000; Vollmer et al., 2017), there have been no direct measurements of corticofugal system function in humans. Theoretically, efferent control of brainstem activity should occur for behaviorally relevant stimuli (Suga, 2008), in states of goal-directed attention (Slee and David, 2015; Vollmer et al., 2017), and strengthen in more taxing listening conditions (e.g., SIN tasks) (Ahissar and Hochstein, 2004). Attention-based neuronal feedback provided by the descending hearing system could help refine early sensory speech representations and assist the brain in navigating complex listening environments.

Here, we aimed to fill two critical voids in our understanding of human auditory system. First, we aimed to determine the degree to which attention actively reinforces early neural representations for speech prior to the neocortex. To this end, we recorded high-density EEGs as listeners performed rapid SIN tasks varying in attentional demand. To overcome challenges of prior work, we used anatomically constrained source imaging to disentangle speech-evoked responses generated from brainstem (FFR) and cortex (ERP). This approach allowed us to jointly index attentional gain in speech processing at both stages of the hearing pathway and evaluate hierarchical processing in auditory attention. Secondly, we aimed to quantify human corticofugal function in SIN perception. We used functional connectivity to characterize “bottom-up” vs. “top-down” (corticofugal) signaling within the brainstem-cortical pathway. Under the premise that the corticofugal system shapes brainstem signal processing only for perceptually taxing and behaviorally-relevant sounds (i.e., “ego-centric selection,” Ahissar and Hochstein, 2004; Suga et al., 2000), we predicted stronger PAC→BS connectivity for noise-degraded relative to clear conditions. Furthermore, if speech perception conforms to early “sensory gating” and “bottleneck” models of attentional locus (Broadbent, 1971; Hirschhorn and Michie, 1990; Picton et al., 1971; Woods and Hillyard, 1978), we hypothesized attention would enhance brainstem phase-locking to speech as measured via source-resolved FFRs (with similar or stronger gains in cortical ERPs). Our results provide the first direct evidence that attention reinforces brainstem auditory processing in humans via corticofugal signaling and confirm this feedback is important for cocktail party speech perception.

## 2. Materials and Methods

### 2.1 Participants

Twenty young adults (age: 18-35 years, *M* = 24, *SD* = 3.4 years; 11 female) participated in the study. An a priori power analysis (t-test, 2-tailed, α = 0.05, power = 95%) revealed this sample was sufficient to detect similar sized effects (*d* = 0.84, 1.0) as in previous FFR/ERP SIN studies (Bidelman et al., 2019; GPower v3.1). All participants exhibited normal hearing thresholds (≤ 25 dB HL; 250-8000 Hz). Because language background and music experience influence FFRs/ERPs and SIN performance (Mankel and Bidelman, 2018; Parbery-Clark et al., 2009; Zhao and Kuhl, 2018), we required participants have < 3 years of formal musical training (*M* = 0.8 years, *SD* = 1.2) and be native English speakers. They were predominantly right-handed (*M* = 82.04%, *SD* = 21.04) (Oldfield, 1971) with no history of neuropsychiatric disorders. All provided written informed consent prior in accordance with protocols approved by the University of Memphis IRB.

### 2.2 Speech stimuli & task

We recorded FFRs and ERPs simultaneously (Bidelman et al., 2013; Bidelman et al., 2019) during SIN-listening tasks designed to evaluate attentional effects on the brain’s hierarchical encoding of speech. Three synthesized vowel tokens (e.g., /a/, /i/, /u/) were presented during the recording of EEGs. We chose vowels as sustained periodic sounds optimally evoke FFRs and ERPs (Bidelman et al., 2019; Skoe and Kraus, 2010a). Each vowel was 100 ms with a common voice fundamental frequency (F0=150 Hz). This F0 is above the phase-locking limit of cortical neurons and observable FFRs in cortex (Bidelman, 2018; Brugge et al., 2009), ensuring our FFRs would be of brainstem origin (Bidelman, 2018; Coffey et al., 2016). Tokens were matched in average r.m.s. amplitude. We presented the vowels in clean (i.e., no background noise) and noise-degraded conditions. For the noise condition, speech stimuli were mixed with 8 talker noise babble (cf. Killion et al., 2004) at a signal-to-noise ratio (SNR) of 5 dB (speech at 75 dB_A_ SPL and noise at 70 dB_A_ SPL). In each SNR, frequent tokens (/a/, /i/) were presented 4000 times while the infrequent token /u/ was presented 140 times (random order; jittered interstimulus = 95-155 ms, 5 ms steps, uniform distribution). Stimulus presentation was controlled by MATLAB (The Mathworks, Inc.; Natick, MA) routed to a TDT RP2 interface (Tucker-Davis Technologies; Alachua, FL) and delivered binaurally through electromagnetically shielded insert earphones (ER-3; Etymotic Research; Elk Grove Village, IL).

Attention was varied via active vs. passive listening blocks. During active blocks, participants had to detect infrequent /u/ tokens via button press. We defined a “hit” as detection within 5 tokens (~500 ms) of a target. For passive blocks, they watched a captioned movie and were instructed to ignore any sounds they heard. We included a control block (*n* = 8) of visual-only stimulation to rule out potential visual confounds inherent to the passive condition. In these runs, the headphone transducers were unplugged from the wall and participants watched the captioned movie (i.e., as in passive block but muted audio). The success of these controls was confirmed by the absence of any time-locked response in the control condition [t-test against FFR_F0_= 0; *t*_83_ = 0.96, *p* = 0.34]), indicating no visual artifacts in the recordings (see Fig. 2). Block presentation (active/passive/control for clean/noisy speech) was counterbalanced across participants to minimize order effects.

### 2.3 QuickSIN test

The Quick Speech-in-Noise (QuickSIN) test assessed listeners’ speech reception thresholds in noise (Killion et al., 2004). Listeners heard lists of 6 sentences, each with 5 target keywords spoken by a female talker embedded in four-talker babble noise. We presented target sentences at 70 dB SPL (binaurally) at SNRs decreasing in 5 dB steps from 25 dB (very easy) to 0 dB (very difficult). We calculated SNR-loss scores as the SNR for 50% keyword recall (Killion et al., 2004). Higher scores indicate poorer SIN performance. We averaged scores from two lists per listener (Fig. 1C).

**Figure 1.**
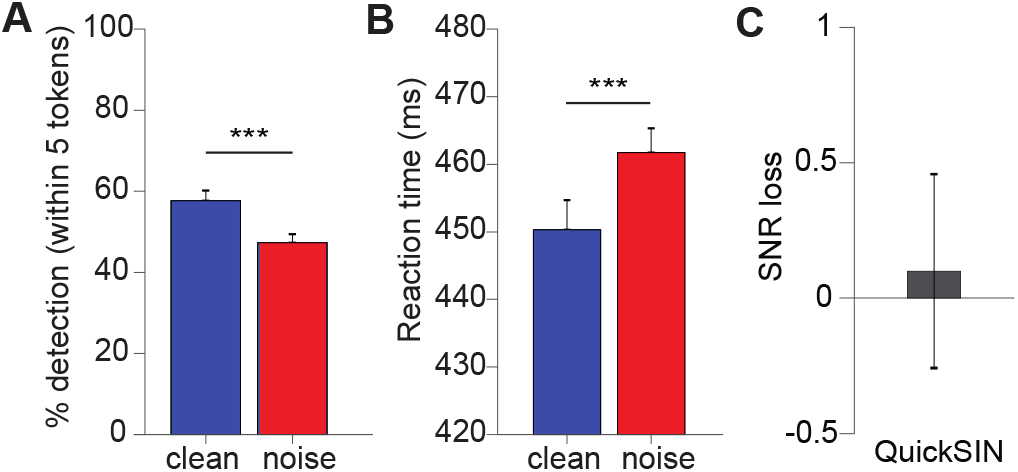
Behavioral target speech detection is hindered by noise. (**A**) Behavioral accuracy and (**B**) reaction times for detecting infrequent /u/ tokens in clean and noise-degraded conditions. Noise hinders speech perception by reducing perceptual accuracy and slowing decision speeds. **(C)** Average QuickSIN scores across participants. errorbars = ± s.e.m., \*\*\**p* < 0.001.

**Figure 2.**
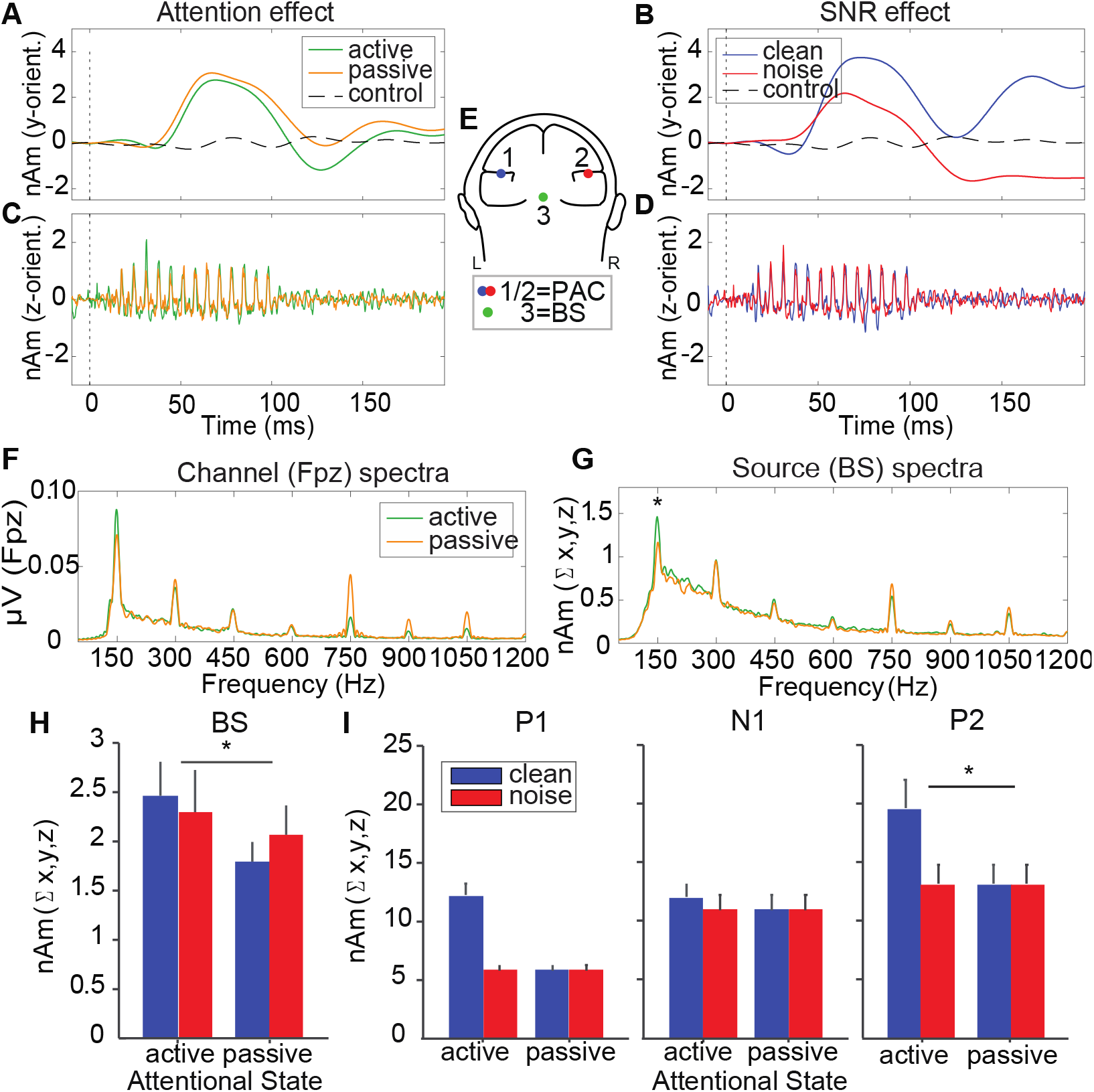
Attention enhances brainstem (FFR) and cortical (ERP) encoding of noise-degraded speech. (**A-B**) Cortical ERPs and (**C-D**) brainstem FFRs extracted from dipoles located in midbrain and bilateral PAC (**E**, inset brain). For clarity, only one dipole orientation is shown per ROI (collapsing tokens). Noise exerts larger effects on cortical vs. brainstem speech coding. Attention enhances both ERPs (P2; 150 ms) and FFRs. (**F-G)** FFR spectra (collapsing tokens/SNRs) illustrating attentional enhancements of speech-FFRs on source (but not channel-level) data. (**H**) Source FFR-F0 amplitudes are stronger during active listening but are resistant to noise. (**I**) Source ERP magnitudes. Attention increases only P2 magnitudes. errorbars = ± s.e.m, \**p* < 0.05. BS, brainstem; PAC, primary auditory cortex.

### 2.4 Electrophysiological recording and analysis

#### 2.4.1 EEG acquisition and preprocessing

We recorded EEGs from 64-channels at 10-10 electrode locations across the scalp (Oostenveld and Praamstra, 2001). Electrodes on the outer canthi and superior/inferior orbit monitored ocular artifacts. Impedances were ≤ 5 kΩ. EEGs were digitized at a high sample rate (5 kHz; DC—2000 Hz online filters; SynAmps RT amplifiers; Compumedics Neuroscan; Charlotte, NC) to recover both fast (FFR) and slow (ERP) frequency components of the compound speech-evoked potential (Bidelman et al., 2013; Musacchia et al., 2008). We processed EEG data in Curry 7 (Compumedics Neuroscan) and BESA Research v7.0 (BESA, GmbH). Ocular artifacts (saccades and blinks) were corrected in continuous EEGs using principal component analysis (PCA) (Picton et al., 2000). Cleaned EEGs were epoched (−10-200 ms), pre-stimulus baselined, and ensemble averaged to obtain compound speech-evoked potentials (Bidelman et al., 2013). We bandpass filtered full-band responses from 130-1500Hz and 1-30 Hz to isolate FFRs and ERPs, respectively (Bidelman et al., 2013; Musacchia et al., 2008). Data were re-referenced to the common average. We excluded infrequent /u/ tokens from the analyses due to their limited number of trials and to avoid mismatch negativities in the data.

#### 2.4.2 Brainstem FFRs

We analyzed the steady-state portion (10-100 ms) of FFR waveforms using a Fast Fourier transform (FFT) which captured the spectral composition of the response. F0 amplitude was quantified as the maximum FFT amplitude within a 10 Hz bin centered around 150 Hz (i.e., F0 of the stimuli). FFR F0 indexes voice pitch coding and predicts successful SIN perception (Mankel and Bidelman, 2018; Parbery-Clark et al., 2009). We analyzed FFRs at both the electrode (Fpz; linked mastoid reference) and source level to compare our findings to previous literature investigating attentional effects on FFRs.

#### 2.4.3 Cortical ERPs

We quantified ERP wave (i.e., P1, N1, P2) amplitude and latency using automated peak analysis. Latency windows were determined following visual inspection of grand average traces. P1 was identified as the maximum positive deflection occurring within 40-80 ms; N1 as the greatest negative deflection between 90-145 ms; and P2 as the maximum positive deflection within 145-175 ms (Hall, 1992).

#### 2.4.4 Source analysis

We transformed listeners’ scalp potentials to source space using BESA which allowed for connectivity analysis between BS and PAC. We used a virtual source montage (Fig. 2E) comprised of regional dipoles (i.e., current flow in x, y, z planes) positioned in the midbrain and bilateral PAC (Bidelman, 2018). This applied an optimized spatial filter to all electrodes that calculated their weighted contribution to the scalp-recorded FFRs/ERPs in order to estimate activity within each source location within the head. For each dipole source, we combined activity from the three orientations (i.e., L2-norm of x, y, z waveforms) to yield an unbiased measure of the aggregate response within each ROI (Coffey et al., 2017). Dipole locations were fixed across subjects. Across stimulus conditions, average goodness of fit (GoF) for our 3-dipole model was 90.6% [residual variance (RV) = 9.4 ± 1.6%], indicating excellent fit to the scalp data.

### 2.5 MRI scans and EEG co-registration

3D T1-weighted anatomical volumes were obtained on a Siemens 1.5T Symphony TIM scanner (tfl3d1 GR/IR sequence; TR=2000 ms, TE=3.26 ms, inversion time=900 ms, phase encoding steps=341, flip angle=8°, FOV=256×256 acquisition matrix, 1.0 mm axial slices). Scanning was conducted at the Semmes Murphey Neurology Clinic (Memphis, TN). Scans were segmented in BESA MRI 2.0. Following inhomogeneity correction (Scherg et al., 2002), images were automatically partitioned into scalp, skull, CSF, and brain compartments (Chan and Vese, 2001) and the cortical surface was reconstructed to allow optional inflation of the brain volume (Fischl et al., 1999). MRI volumes were rendered in both ACPC and Talairach (Talairach and Tournoux, 1988) spaces using 3D spline interpolation.

Following MRI segmentation, electrode locations were warped to the scalp surface (anchored to the nasion and preauricular fiducials) to co-register sensor locations to individuals’ anatomy. Electrode positions were mapped with a quad sensor Polhemus Fastrak digitizer (Polhemus, Colchester, VT). We then generated a 4-layer finite element head model (FEM) based on the MRI segmentation (Wolters et al., 2007) to construct each individuals’ leadfield (forward volume conductor). The FEM leadfield described the magnitude each source signal contributed at each sensor (Scherg, 1990) and is less prone to spatial errors than other head models (e.g., concentric spherical conductor) (Fuchs et al., 2002). Collectively, this approach allowed us to source localize each participant’s cortical ERPs and brainstem FFRs with high precision, constrained to their individual brain anatomy. MRIs were not available for 4 participants. In these cases, we used a 4-shell spherical volume conductor head model and an adult template anatomy (Berg and Scherg, 1994).

### 2.6 Functional connectivity

We measured functional connectivity between PAC and BS source waveforms using phase transfer entropy (PTE), a measure of nonlinear, directed (causal) signal dependency (Bidelman et al., 2018; Bidelman et al., 2019; Price et al., 2019). For details, see Lobier et al., 2014. PTE was computed in both directions to quantify differences in the strength of bottom-up afferent (BS→PAC) vs. top-down efferent (PAC→BS) connectivity within the auditory brainstem-cortical pathway. This allowed us to evaluate how attending to auditory signals and listening in more adverse conditions influenced bidirectional signaling including critical corticofugal (PAC→BS) function.

### 2.7 Statistical analyses

We performed 2×2×2 (vowel × attention × SNR) mixed model (subjects=random factor) ANOVAs (GLIMMIX, SAS^®^ 9.4, SAS Institute; Cary, NC). Tukey-Kramer adjustments corrected multiple comparisons. Data were log-transformed to satisfy normality and homogeneity of variance assumptions. Paired samples *t*-tests (two-tailed) were used to compare behavioral performance between conditions. A generalized linear model (GLM) evaluated brain-behavior relationships. Neural measures from each ROI (FFR_BS_: F0 amplitude; ERP_PAC_: P2 magnitude), connectivity (afferent, efferent), behavioral [QuickSIN, pure-tone average (PTA) hearing thresholds]), and stimulus factors (SNR) were included as predictors of perceptual throughput for the active SIN detection task [throughput ~ 1 + FFR + ERP + aff + eff + QSIN + PTA + SNR]. Behavioral throughput reflects the time-accuracy tradeoff (i.e., throughput = % / RT for target detection) whereby slower response speeds result in poorer throughput and thus less overall perceptual efficiency (Bidelman et al., 2014; Salthouse and Hedden, 2002).

### 2.8 Data availability

The data supporting the reported findings are available from the corresponding author upon reasonable request.

## 3. Results

### 3.1 Behavioral data

Noise expectedly reduced listeners’ target speech detection accuracy (*t*_19_ = 4.48, *p* < 0.001) and slowed response speeds (*t*_19_ = −4.78, *p* < 0.001), confirming poorer SNRs were detrimental to speech perception (Fig. 1A, B).

### 3.2 Electrophysiological data

Source-level FFRs and ERPs reflecting brainstem and cortical activity contrast the effects of attentional state (Fig. 2A, C) and noise (Fig. 2B, D) on the neural encoding of speech during the perceptual task.

#### 3.2.1 Brainstem FFRs

To determine if attention modulates brainstem SIN processing, we measured FFR F0 amplitudes, a neural proxy of voice pitch encoding that serves as an important cue for tracking speech in background noise (Assmann, 1996; Mankel and Bidelman, 2018). We analyzed FFRs at both electrode and source levels to replicate previous literature. Noise weakened scalp FFRs (*F*_1, 136_ = 8.15, *p* = 0.01) as in previous studies. Critically, we found no evidence for attentional effects in scalp (electrode-level) data (Fig. 2F; *F*_1, 136_ = 1.68, *p* = 0.20), corroborating prior null attention effects on FFRs (Picton and Hillyard, 1974; Picton et al., 1971; Woods and Hillyard, 1978). Contrastively, *source-level* FFRs were enhanced during active listening (Fig 2G, H; *F*_1, 136_ = 5.39, *p* = 0.02) and were invariant to noise (*F*_1, 136_ = 2.22, *p* = 0.14). These findings reveal attention modulates brainstem speech encoding but only when viewed at the source level.

#### 3.2.2 Cortical ERPs

For cortical source ERPs (i.e., activity from PAC), pooling hemispheres, P1 and P2 magnitudes decreased with noise (P1: *F*_1, 136_ = 77.33, *p* < 0.0001; P2: *F*_1, 136_ = 6.21, *p* = 0.01), whereas N1 differences were negligible (*F*_1, 136_ = 0.04, *p* = 0.84) (Fig. 2I). Critically, attention enhanced P2 (*F*_1, 136_ = 4.97, *p* = 0.03), confirming its role in active listening (Naatanen, 1975; Picton and Hillyard, 1974) and a biomarker of SIN perception (Bidelman et al., 2019). No other significant effects or interactions were found.

### 3.3 Brainstem-cortical functional connectivity

We used phase-transfer entropy (PTE) to quantify directional functional connectivity and evaluate effects of attention and task demands on bottom-up (i.e., afferent) and top-down (i.e., efferent) neural signaling within the brainstem-cortical pathway (see Section 2.6). Connectivity was stronger during active vs. passive listening overall (Fig. 3A, C; attention main effect: *F*_1, 295_ = 7.92, *p* = 0.01). More critically, we found a direction × SNR interaction on connectivity strength (Fig. 3B, C; *F*_1, 295_ = 11.59, *p* < 0.001). Efferent (PAC→BS) connectivity was more robust for clean speech and weakened in noise (*t*_295_ = 3.32, *p* = 0.01), whereas afferent (BS→PAC) connectivity was invariant to noise (*t*_295_ = −1.50, *p* = 0.44). Additional planned contrasts revealed noise decreased efferent connectivity during passive (*t*_292_ = 3.12, *p* = 0.04) but not active SIN perception (*t*_292_ = 1.57, *p* = 0.77). These findings reveal corticofugal transmission remained stable during active SIN perception but disengaged while passively coding otherwise identical speech stimuli.

**Figure 3.**
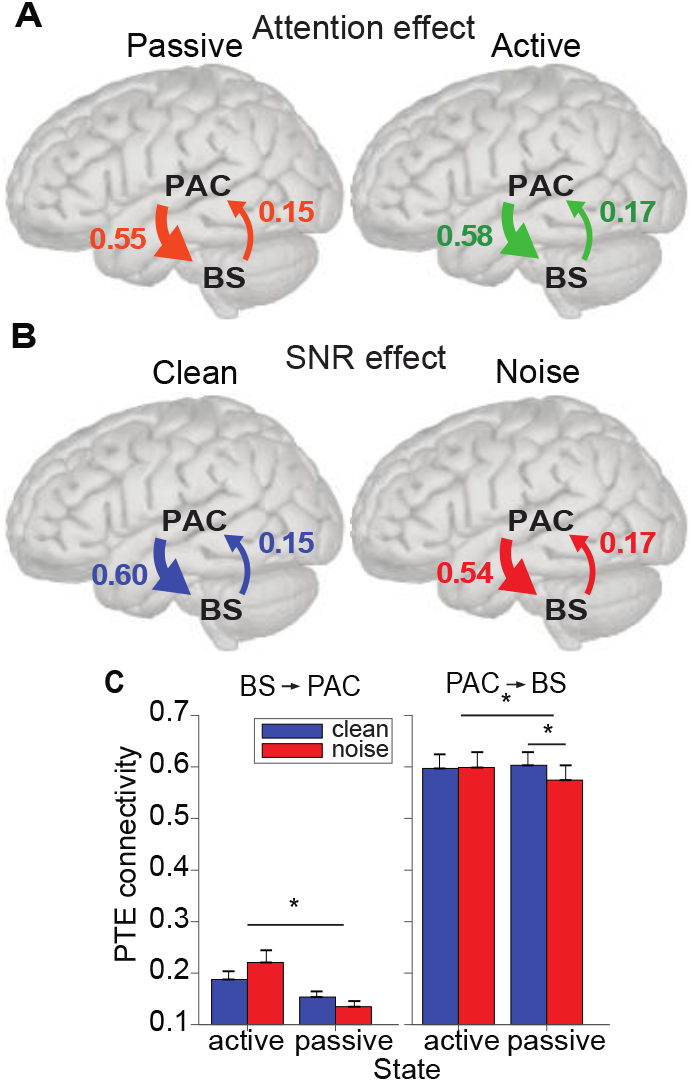
Attention and noise modulate afferent and efferent connectivity between brainstem and cortex during speech perception. Functional connectivity measured via phase-transfer entropy between BS and PAC source waveforms. Main effect of (**A**) attention and (**B**) SNR on connectivity strength. (**C**) Attention increases bidirectional communication between BS and PAC. Corticofugal efferent connectivity decreases in noise, but only during passive listening. Attention maintains efferent signaling in more challenging listening conditions. errorbars = ± s.e.m, \**p* < 0.05.

### 3.4 Brain-behavior correlations

Having established attentional gain effects within and between BS and PAC, we next used GLM regression to evaluate relations between neural, behavioral, and stimulus factors that drive behavioral SIN processing, as summarized via perceptual throughput (i.e.,% / RT~ 1 + FFR + ERP + aff + eff + QSIN + PTA + SNR) (Bidelman et al., 2014; Salthouse and Hedden, 2002). The overall multivariate model was highly significant (*F*_7,72_ = 5.83, *p* < 0.0001; *r*^2^ = 0.27). Evaluating individual terms (see Inline Supplementary Table 1) revealed significant predictors in F0 amplitude (*t*72 = 2.22, *p* = 0.03), afferent connectivity (*t*_72_ = −2.19, *p* = 0.03), QuickSIN scores (*t*_72_ = −2.51, *p* = 0.01), and SNR (*t*_72_ = −4.39, *p* < 0.0001). These results confirm the behavioral relevance of BS-PAC connectivity as well as brainstem FFRs in predicting successful SIN perception.

## 4. Discussion

By simultaneously recording speech-evoked brainstem and cortical EEGs and manipulating attentional engagement and task difficulty during active SIN perception, our findings show (i) attention actively modulates SIN processing across the auditory neuroaxis, exerting influences on speech representations as early as the midbrain; and (ii) attention reinforces top-down neural communication from PAC to BS in adverse listening conditions. Our neuroimaging results establish a direct role of the human corticofugal feedback system as an aid to cocktail party speech perception.

At the cortical level, we found source ERPs were diminished in noise, confirming acoustic degradation weakens neural representations for speech in PAC (Bidelman et al., 2019; Du et al., 2014). Contrasting cortex, brainstem FFR-F0 was surprisingly resistant to noise. The resilience (and even enhancement) of FFR voice pitch encoding in low-level noise is attributable to a reinforcement of neural activity as midbrain neurons phase-lock to both F0 and upper harmonics of vowel stimuli (Bidelman and Krishnan, 2010; Parbery-Clark et al., 2009; Smith et al., 1978). Collectively, our concurrent subcortical-cortical recordings expose a differential pattern in how noise challenges the brain’s speech processing with stronger changes in cortex relative to brainstem levels.

Our findings corroborate previous neuroimaging studies suggesting both FFRs and the cortical P2 are strong predictors of successful SIN perception in both younger (Bidelman et al., 2018; Coffey et al., 2017) and older (Bidelman et al., 2019) adults. That P2 is more strongly related to SIN perception than earlier ERP waves (e.g., P1, N1) is consistent with prior neuroimaging studies linking P2 to auditory perceptual object formation (Bidelman et al., 2013) and P1/N1 to exogenous acoustic properties (Bidelman et al., 2018; Parbery-Clark et al., 2011). A potential explanation for our cross-level differences is that cortical ERPs reflect aggregate coding of multiple and integrated acoustic features (e.g., pitch, timbre, etc.), whereas FFR-F0 reflects primary voice pitch encoding (Bidelman and Krishnan, 2010; Skoe and Kraus, 2010a). Collectively, our electrophysiological data coupled with prior studies provide converging evidence that noise differentially influences speech coding at brainstem and cortical levels.

We found pervasive enhancements in neural coding with active listening at both subcortical and cortical levels. Our ERP data replicate well-known attentional effects in auditory cortex and its links to complex speech processing (Bidelman et al., 2019; Parbery-Clark et al., 2011). ERP amplitudes typically show attention-driven enhancement of the sensory-perceptual N1 response with associated reductions in P2 (Naatanen, 1975). Even larger attentional gains are observed in late cortical activity indexing post-perceptual processing and response selection (Picton and Hillyard, 1974; Picton et al., 1971). Our ERP data reveal larger P2 magnitudes but invariant N1 across attentional states. These findings are consistent with notions that N1 reflects changes in arousal and acoustic feature coding (Bidelman et al., 2013; Coull, 1998), whereas P2 indexes active perceptual processes including stimulus classification, speech identity, and auditory object formation (Bidelman et al., 2013). The lack of attentional effects on N1 might also be attributable to heavier neural adaptation given the rapid delivery of our speech stimuli. Indeed, PAC attention effects are particularly susceptible to stimulus presentation rate in the timeframe of N1 (~100 ms) (Neelon et al., 2006).

Critically, our data expose strong attentional enhancements in brainstem speech-FFRs when viewed at the *source* level. Attention effects on FFRs have been highly controversial (Dunlop et al., 1965; Galbraith and Kane, 1993; Picton et al., 1971; Varghese et al., 2015); brainstem responses recorded during speech perception tasks typically fail to vary with listening state (Varghese et al., 2015), despite concomitant changes in cortical ERPs. Our data help reconcile equivocal findings by revealing top-down influences in source but not channel-level (i.e., scalp electrode) FFR data. Previous failures to consistently observe attentional changes in FFRs might rest in the overwhelming analysis of scalp-level data which blur activity of multiple generators underlying the FFR including BS and PAC (Bidelman, 2018; Coffey et al., 2016), but also cochlear sources (Bidelman, 2018) that may be too peripheral for the purview of attention. Moreover, the high voice pitch (F0=150 Hz) of our stimuli rules out cortical contributions to our FFRs, which were recorded above the phase-locking limit of PAC neurons (<100 Hz) (Bidelman, 2018). Consequently, our source analysis reveals that “purer” FFRs localized to brainstem (cf. Hartmann and Weisz, 2019) are highly sensitive to attention-dependent reshaping. Our results provide convincing evidence that attention enhances speech coding online as early as the midbrain and suggest subcortical structures function as an early filtering mechanism as posited by early attention theories (Broadbent, 1971; Treisman, 1960).

Beyond local enhancements, our data further show attention strengthens neural signaling in both feedforward (afferent) and feedback (efferent) directions within the primary auditory pathways. In the visual system, attention increases inter-regional connectivity involved in sensory processing (Buchel and Friston, 1997) with attentional selection being driven by interactions between feedforward and feedback mechanisms (Khorsand et al., 2015). In audition, we found top-down feedback is stronger than its feedforward counterpart, surprisingly, regardless of attentional state or task difficulty (low-level noise). Yet, attention enhanced BS-PAC neural communication bidirectionally. Higher engagement of the efferent system regardless of attention may explain why some FFR studies have observed response enhancements even in passive speech listening tasks (Chandrasekaran et al., 2009; Skoe and Kraus, 2010b).

Animal studies show corticofugal efferents tune subcortical auditory signal processing during short-term auditory learning suggesting cortically-guided feedback shapes earlier sensory coding (Bajo et al., 2010; Suga, 2008). Analogous corticofugal modulation is speculated to account for signal enhancements observed in human FFRs among listeners with long-term experience and training (Chandrasekaran et al., 2009; Galbraith et al., 2003; Lukas, 1980; Skoe and Kraus, 2010b; Tzounopoulos and Kraus, 2009; Wong et al., 2007). To date, evidence for direct corticofugal involvement in human hearing has been unverified. Theoretically, increased top-down contributions are expected in more challenging scenarios (e.g., during learning, degraded listening environments, increased attentional demands) to sharpen earlier sensory processing and facilitate transmission of faithful neural representations of the acoustic input. Using direct measures of brainstem-cortical connectivity, we find attention reinforces not only brainstem speech coding but also corticofugal signaling in more difficult (noisy) conditions as evidenced by weakened efferent connectivity for passive listening in noise yet sustained connectivity for active listening. Our findings suggest attention maintains top-down (and bottom-up) neural signaling in noise to tune and enhance early speech encoding which is associated with improved behavioral performance.

While our data show clear attentional modulation of brainstem speech processing, they cannot adjudicate the domain generality of these effects. It remains possible that similar attentional benefits in the cortico-collicular system exist for non-speech sounds, as suggested in animal data (Suga et al., 2000; Suga and Ma, 2003). Additionally, our analyses were restricted to early portions of the auditory-speech network (i.e., BS↔PAC) that reflect high-fidelity sensory coding but do not encompass the later neural networks implicated in semantic, lexical, and other post-perceptual processing necessary for spoken word recognition (e.g., frontoparietal cortices). Interhemispheric connections contribute to auditory attention (Bamiou et al., 2007; Petersen and Posner, 2012), and activation within secondary auditory and prefrontal regions (i.e., posterior superior temporal gyrus, inferior frontal gyrus, prefrontal cortex) strengthens during demanding speech perception tasks (Hickok and Poeppel, 2000). Importantly, we do not deny the very necessary contributions of higher-order brain structures and related attentional networks that engage in challenging listening conditions to decode speech (Alain et al., 2018; Du et al., 2014). Rather, our data argue these processes commence much earlier in the speech hierarchy with attention tuning the representation and formation of speech percepts prior to cortex.

## 5. Conclusions

In sum, our results emphasize the complex interaction between attention and hierarchical auditory processing during SIN perception. Our results provide evidence to resolve ongoing debates regarding attentional influences on early brainstem encoding and corticofugal engagement during active listening in humans. We show attention enhances neural encoding in cortex but also in brainstem, a surprisingly early stage of processing. Furthermore, using functional connectivity, we demonstrate attentional enhancement in two-way communication between subcortical and cortical levels with stronger communication occurring in the corticofugal direction. Thus, our results suggest attention serves as a mechanism to overcome detrimental noise effects and maintain efficient top-down signaling in challenging listening conditions. Overall, our findings provide novel measurements of corticofugal function in human hearing and establish its involvement as an attention-dependent gain control in speech perception.

## Acknowledgements

This work was supported by the National Institutes of Health (NIH/NIDCD R01DC016267) (G.M.B.) and the UofM Institute for Intelligent Systems Dissertation grant (C.N.P.).

