## Supplemental Table 1 for "Attention reinforces human corticofugal system to aid speech perception in noise"

**Table S1.** GLME model fit parameters for prediction of behavioral throughput. Coefficients and significance tests for individual predictor variables including neural and hearing measures and SNR. Level of significance denoted *$p < 0.05$, ***$p < 0.001$

| Name | Estimate | SE | tStat | DF | pValue | Lower | Upper |
|---|---|---|---|---|---|---|---|
| Intercept | 0.1633 | 0.0197 | 8.29 | 72 | 4.41e-12 | 0.1240 | 0.2026 |
| FFR* | 0.0044 | 0.0020 | 2.22 | 72 | 0.03 | 0.0005 | 0.0084 |
| ERP | -0.0004 | 0.0003 | -1.55 | 72 | 0.13 | -0.0009 | 0.0001 |
| aff* | -0.0793 | 0.0363 | -2.19 | 72 | 0.03 | -0.1517 | -0.0070 |
| eff | 0.0043 | 0.0241 | 0.18 | 72 | 0.86 | -0.0437 | 0.0524 |
| QSIN* | -0.0043 | 0.0017 | -2.51 | 72 | 0.01 | -0.0077 | -0.0009 |
| PTA | -0.0003 | 0.0007 | -0.47 | 72 | 0.64 | -0.0018 | 0.0011 |
| SNR*** | -0.0237 | 0.0054 | -4.39 | 72 | 3.87e-05 | -0.0345 | -0.0130 |
